# Where do I go? Decoding temporal neural dynamics of scene processing and visuospatial memory interactions using CNNs

**DOI:** 10.1101/2024.12.17.628860

**Authors:** Clément Naveilhan, Raphaël Zory, Stephen Ramanoël

**Author notes:** CA, Corresponding Author: Clément Naveilhan.

## Abstract

Visual scene perception enables rapid interpretation of the surrounding environment by integrating multiple visual features related to task demands and context, which is essential for goal-directed behavior. In the present work, we investigated the temporal neural dynamics underlying the interaction between the processing of visual features (*i*.*e*., bottom-up processes) and contextual knowledge (*i*.*e*., top-down processes) during scene perception. We analyzed EEG data from 30 participants performing scene memory and visuospatial memory tasks in which we manipulated the number of navigational affordances available (*i*.*e*., the number of open doors) while controlling for similar low-level visual features across tasks. We used convolutional neural networks (CNN) coupled with gradient-weighted class activation mapping (Grad-CAM) to assess the main channels and time points underlying neural processing for each task. We found that early occipitoparietal activity (50-250 ms post-stimulus) contributed most to the classification of several aspects of visual perception, including scene color, navigational affordances, and spatial memory content. In addition, we showed that the CNN successfully trained to detect affordances during scene perception was unable to detect the same affordances in the spatial memory task after learning, whereas a similarly trained and tested model for detecting wall color was able to generalize across tasks. Taken together, these results reveal an early common window of integration for scene and visuospatial memory information, with a specific and immediate influence of newly acquired spatial knowledge on early neural correlates of scene perception.

## Introduction

Visual scene perception is a critical cognitive function that enables individuals to rapidly encode, interpret, and interact with their environment. This process requires the integration of low-, mid-, and high-level visual information (*e*.*g*., object features, spatial configurations, and landmarks; Bartnik & Groen, 2023) to support efficient and adaptive behavior in dynamic settings such as natural environments (Epstein & Baker, 2019). The complexity of visual perception becomes even more pronounced in everyday life when we consider the variety of goals, tasks, contexts, or prior knowledge that modulate scene processing and its neural correlates (Bar, 2009; Bar et al., 2006; Kay et al., 2023; Malcolm et al., 2016; Nau et al., 2024; Ritchie et al., 2024). Therefore, a deeper understanding of the neural dynamics underlying scene perception, especially the interaction between bottom-up (*e*.*g*., visual features) and top-down (*e*.*g*., prior knowledge) information, is essential for elucidating the mechanisms of visual cognition.

In this context, the processing of navigational affordances available in visual scenes represents a promising theoretical and methodological framework (Bartnik et al., 2024; Djebbara et al., 2019; Gregorians & Spiers, 2022; Naveilhan et al., 2024). Initially proposed to be automatically extracted solely by bottom-up mechanisms during scene perception (Bonner & Epstein, 2017; Harel et al., 2022), recent findings also suggest the influence of contextual information on scene processing at both behavioral and neural levels (Aminoff & Tarr, 2021; Choi et al., 2020; Naveilhan et al., 2024). For example, Naveilhan et al. (2024) reported that increasing the number of navigational affordances (*i*.*e*., open doors in a room) selectively decreased participants’ accuracy in retrieving the position of a previously learned goal. These behavioral findings were complemented by neural data using event-related potentials (ERPs) recorded from occipitoparietal electrodes spanning scene-selective regions (SSRs), specifically the occipital place area (OPA), which plays a critical role in encoding pathways for movement in scenes (Bonner & Epstein, 2017; Harel et al., 2016, 2022; Kaiser et al., 2020; Kamps et al., 2024). These findings suggest that prior spatial knowledge modulates early neural markers of visual processing, specifically the P2 component (Harel et al., 2016, 2022; Kaiser et al., 2020). However, the restriction of previous analyses to ERPs over occipitoparietal electrodes may have limited the ability to investigate the complex global dynamics of visual information processing involved in visuospatial tasks (Groen et al., 2017; Kravitz et al., 2011).

Recent advances in signal processing and supervised learning with neural networks now make it possible to consider multiple brain regions that support complex dynamics for visual perception. Notably, Orima and Motoyoshi (2023) trained a convolutional neural network (EEGNet; Lawhern et al., 2018) to classify both scene categories and their global properties (*e*.*g*., naturalness, openness, roughness) based on EEG data. By applying gradient-weighted class activation mapping (Grad-CAM; Selvaraju et al., 2017), the authors reported that early EEG signals from occipital electrodes were crucial for classifying openness, whereas later signals from frontal electrodes were crucial for determining naturalness and scene categories. These findings suggest that different properties of visual scenes are processed in different brain regions and at different times. Dwivedi et al. (2024) further explored this complex sequencing of scene perception using representational similarity analysis to compare neural responses across occipital electrodes with computational models of 2D, 3D, semantic features, and navigational affordances. Their results showed that visual features are processed earlier in the temporal sequence (130-170 ms post-stimulus) than navigational affordances (around 300 ms), suggesting hierarchical processing in line with the findings of the Orima group. However, these results also seem to be partially inconsistent with those of Harel et al. (2022), who showed that navigational affordances were processed earlier (around 230 ms). A possible explanation is based on the fact that in previous protocols participants were passively presented with the scene (Bonner & Epstein, 2017; Harel et al., 2022), whereas in Dwivedi et al. (2024) participants had explicitly engaged in navigational affordance processing. These differences in task engagement, contextual information, and the visual features of the stimuli used could influence the temporal dynamics of navigational affordance extraction, as previously reported only at the behavioral level (Naveilhan et al., 2024). However, the temporal dynamics of the neural correlates underlying the interaction between scene information processing and prior contextual information remain elusive and represent a major challenge for current research in visual cognition (Nau et al., 2024; Ritchie et al., 2024).

To address this issue, we performed novel analyses using a convolutional neural network (CNN) on a previously acquired dataset, obtained during scene perception and visual spatial memory tasks in which we manipulated the number of available affordances while controlling for similar low-level visual features across tasks (Naveilhan et al., 2024). We included all 64 available electrodes to investigate the global dynamics of the expected interaction between task-relevant top-down (*i*.*e*., prior spatial knowledge) and bottom-up information processing (*i*.*e*., visual features). The aim of the present study is to provide new insights into the temporal neural dynamics underlying task-specific visuospatial representations by leveraging the complexity of CNNs to disentangle the intricate structure of EEG data. Specifically, based on previous work on visual object processing for action (Enge et al., 2023), we hypothesize that top-down information modulates the early neural signature of scene perception, including neural correlates of visual scene features. In this sense, we expect that processing of prior spatial knowledge will modulate early occipitoparietal activity (around 200 ms), allowing the CNN to accurately decode the task participants are performing. We also propose that this early time window contains specific information about the previously learned goal position. Furthermore, we hypothesize that learned spatial information will modulate the early neural signature of affordance processing despite identical visual features. In other words, a CNN trained to decode the navigational affordances available when participants have no prior spatial knowledge will not be able to decode the affordance information after participants have learned the goal position, even when the visual scenes are the same.

## Method

### Participants

In the present study, we conducted a reanalysis of a dataset from a previously published study (Naveilhan et al., 2024). Electroencephalographic (EEG) data were collected from 30 young adults (mean age = 24.31 years, SD = 0.65, range = 19-31; 16 females). Sample size was determined a priori using G*Power (version 3.1.9.7; Faul et al., 2007) based on a previous EEG study of scene perception (Naveilhan et al., 2023), which indicated that a minimum of 23 participants was required to achieve statistical power of 0.95 at an alpha level of 0.05. The study was approved by the local ethics committee (CERNI-UCA opinion no. 2021-050), and all participants gave informed consent prior to participation. They were all right-handed, had normal or corrected-to-normal vision, and had no history of neurological or cognitive disorders.

### Stimuli and procedure

The experiment used visual stimuli developed with Unity Engine (v2019.2.0.0f1) and presented on an iiyama ProLite B2791HSU monitor (1920×1080 resolution, 30-83 Hz) positioned 60 cm from the participants. Stimulus presentation was controlled by PsychoPy (v2022.13) running on a Dell Precision 7560 workstation with an Intel Xeon W-11955 processor. The stimuli consisted of images of simple rectangular rooms, each containing either a door or a gray rectangle on three visible walls. These designs were adapted from previous studies (Bonner & Epstein, 2017; Harel et al., 2022). To avoid potential associations between door locations and wall colors, seven different door configurations and seven wall color variations were used, resulting in 49 unique stimuli. All stimuli are publicly available in the OSF repository, and a subset is presented in **Figure 1**.

**Figure 1.**
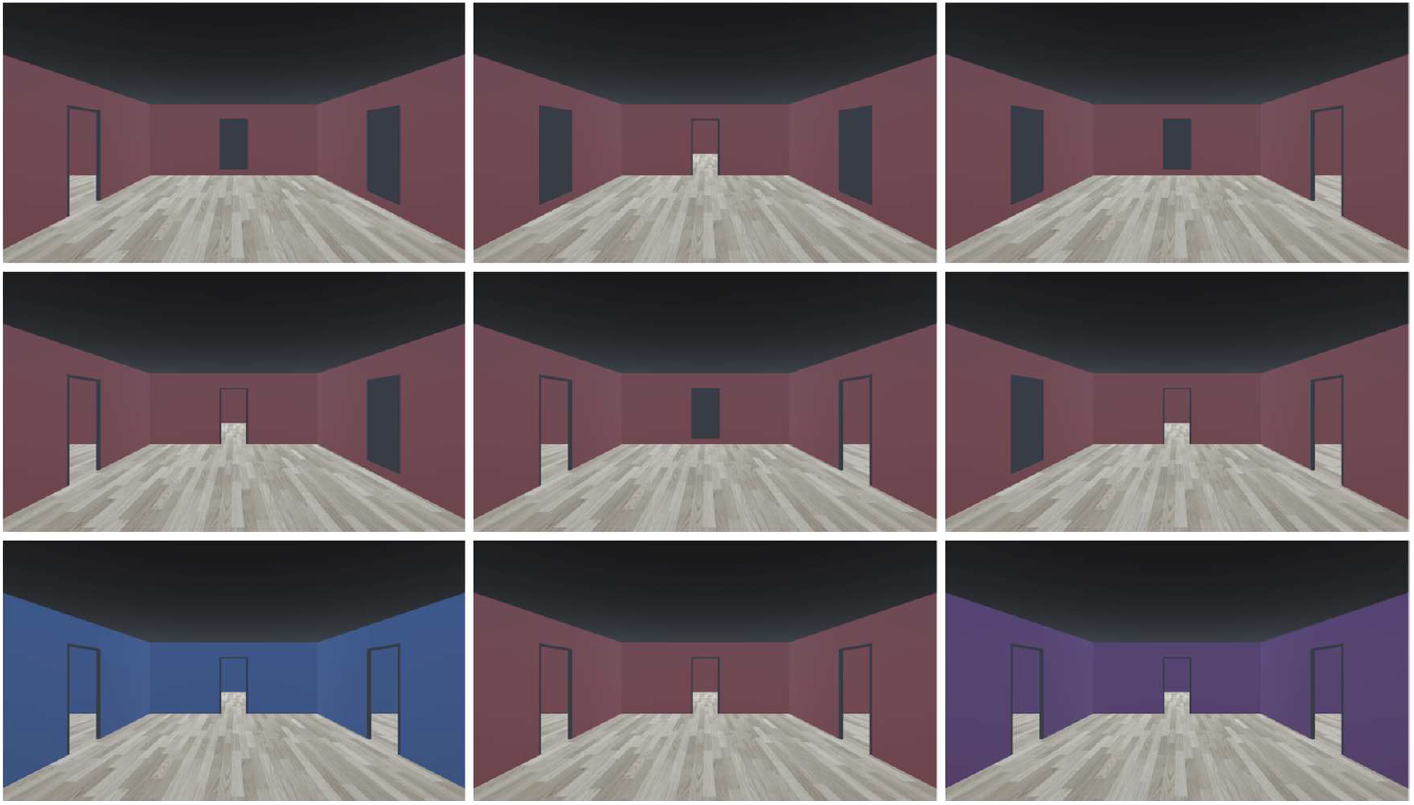
Presentation of a subset of the stimuli used, showing the 7 possible affordance configurations and 3 of the 8 wall colors included in the protocol. The complete set of stimuli is available in the OSF repository.

Participants completed two tasks: a scene memory task and a spatial memory task, each organized into six randomized blocks. Within each block, the order of stimulus presentation was pseudorandomized to maintain participant engagement. In the scene memory task, participants performed a 1-back task in which they identified whether the current scene matched the previous scene based on wall color only, pressing a designated key upon detection of a match. This task consisted of 64 images with four door configurations presented in two different environments, each repeated eight times. The spatial memory task required participants to first passively navigate through rooms to memorize the path to a hidden goal that could be located to the left, center, or right. Each wall color corresponded to a specific goal direction. Participants were then presented with the images and had to indicate the direction of the goal by pressing a key on a USB keyboard as quickly and accurately as possible. Although the same images were used as in the scene memory task, participants now had access to spatial contextual information because they knew that a goal was hidden behind one of the doors. The spatial memory task consistently followed the scene memory task within each block to prevent prior knowledge from influencing performance during the scene memory phase.

For each task, images were displayed for 1 second, followed by a fixation cross lasting between 0.5 and 1 second, and auditory feedback for responses. Over the course of the experiment, participants viewed a total of 1,536 images (768 per task), and each session lasted approximately 55 minutes.

### EEG preprocessing

EEG data were acquired at a sampling rate of 500 Hz using a 64-channel Waveguard™ cap with active wet electrodes, connected to an eego™ mylab amplifier (ANT Neuro) and digitized at 24-bit resolution. The reference electrode was positioned at CPz, and the ground electrode at AFz. Electrode impedances were maintained below 10 kΩ, with the majority under 5 kΩ. EEG recordings were synchronized with stimulus presentations using LabRecorder in the Lab Streaming Layer (Kothe et al., 2024).

Since preprocessing steps have been proposed to play an important role in the decoding capabilities of EEGNet (Kessler et al., 2024), we opted for an open source and fully reproducible pipeline implemented in MATLAB (R2024a), namely, the BeMobil pipeline (Klug et al., 2022) using EEGLAB v2024.2 (Delorme & Makeig, 2004). Non-experimental segments and highly artifacted portions of the data were manually excluded and the signals were downsampled to 250 Hz. Line noise was automatically detected and removed using Zapline Plus (Klug & Kloosterman, 2022). Bad electrodes were detected based on a correlation threshold of 0.8 and a maximum downtime of 0.6, and then interpolated using spherical interpolation methods, with an average of 3.10 ± 2.58 electrodes removed per subject. The data were referenced to the common average. Additional artifact cleaning was performed in the time domain using Artifact Subspace Reconstruction (ASR) with a cutoff threshold of 20 (Chang et al., 2020), resulting in the removal of an average of 7.51 ± 3.52% of data points.

Subsequently, the data were high-pass filtered at 1.5 Hz and AMICA decomposition was applied with 10 rejection iterations and a sigma threshold of 3 (Klug et al., 2024; Klug & Gramann, 2021). Dipole fitting was performed using DipFit, and independent components were classified using ICLabel with default parameters (Pion-Tonachini et al., 2019). Components identified as muscle, line noise, eye, or heart artifacts were excluded, and only those classified as brain or other with residual variance less than 15% were retained (Delorme et al., 2012). The data were then band-pass filtered between 0.3 and 50 Hz, and epochs from 1 second before to 2 seconds after stimulus onset were extracted. Epochs containing artifacts larger than 100 µV were excluded, resulting in an average of 1,461 ± 90.02 epochs per subject, balanced across conditions.

### EEGNet and Grad-CAM procedure

Data were exported from MATLAB to Python for subsequent analyses using an EEG-based classification framework implemented with PyTorch (Paszke et al., 2019). Analyses were performed on an NVIDIA RTX 4090 GPU with 24 GB of VRAM using CUDA 12.6. Each dataset was normalized and balanced to ensure equal class representation. Our methodology relied on the EEGNet architecture (Lawhern et al., 2018), which incorporates convolutional layers, batch normalization (Ioffe & Szegedy, 2015), exponential linear unit (ELU) activation functions, and optimized dropout layers (Srivastava et al., 2014) to mitigate overfitting. By capturing both spatial patterns across electrodes and temporal dynamics within single trials, EEGNet is well suited to handle the complexity of the present dataset while maintaining computational efficiency for single trial classification. Optuna’s hyperband pruner (Akiba et al., 2019; Li et al., 2018), which uses a tree-structured Parzen estimator, was applied for optimal parameter selection, including learning rate and dropout rate, with early stopping based on validation accuracy. To address class imbalances and improve model generalization, class-weighted cross-entropy loss was used (Cui et al., 2019). After training the model on 80% of the full pooled dataset with early stopping, performance was evaluated using 5-fold cross-validation for each individual subject. Classification accuracy was then compared to chance levels using a binomial cumulative distribution approach. For each analysis, a binomial distribution was computed using MATLAB’s binoinv function, taking into account the predefined significance level (α = 0.05), the number of predictions (the average number of classifications made per participant), and the number of classes (2 for task predictions, 7 for affordances, and 8 for wall color). The binomial method was used to determine statistical significance, as it is comparable in reliability to permutation testing for datasets with more than 100 trials, but does not have the extensive computational requirements of permutation testing (Combrisson & Jerbi, 2015; Vahid et al., 2020). We then used 100,000 iterations of bootstrap resampling to calculate the mean difference between the model accuracy and a random sample drawn from the generated binomial distribution. From the bootstrap distribution of the mean differences between accuracy of the model and chance, we derived a 95% confidence interval and a *p-value*, which are presented in the Results section. Finally, Grad-CAM (Selvaraju et al., 2017) was used to visualize the contributions to classification by focusing on activations from the second convolutional block (Orima & Motoyoshi, 2023). Specifically, gradients of predicted values were computed with respect to feature map activations, resulting in two-dimensional localization maps (electrodes and time points). These maps were processed through ReLU layers, normalized, and averaged across participants to identify the key EEG channels and time points that most significantly contributed to the classifications, as illustrated in the Results section. To ensure the robustness and generalizability of our findings, all analyses were repeated five times, and the median results were reported. Detailed information about the different models architectures and example code are available in the study’s OSF repository. We decided to chose hyperparameter selection using Bayesian optimization and HyperBand (Falkner et al., 2018), which addresses the main limitation of hyperparameter selection highlighted previously (Orima & Motoyoshi, 2023), but still relies on some hyperparameters that may introduce small inconsistencies in test-retest reliability. However, for each analysis, hyperparameter selection, training, and testing were performed five times, and the accuracy of the model for all analyses was consistent across the five repetitions.

## Results

### Early EEG signal is modulated by visuospatial memory information

In the first analysis, we merged the entire dataset, performed hyperparameter optimization and then trained the model on the balanced data to classify the tasks participants were performing (*i*.*e*., scene memory or spatial memory). For each subject, we then performed 5-fold cross-validation and used Grad-CAM on the second layer of the trained CNN (**Figure 2**) to identify the time points and electrodes that contributed most to classifying the task participants were performing (**Figure 2**).

**Figure 2.**
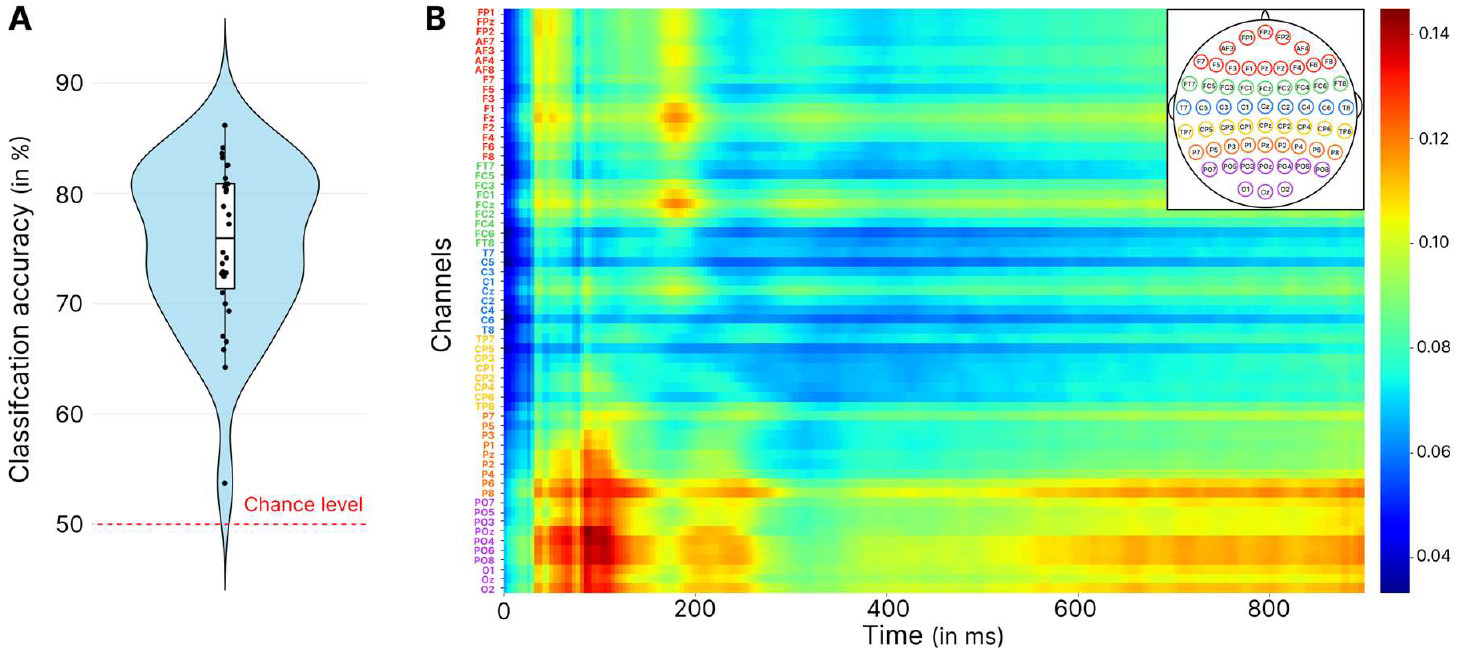
Results of the EEGNet classification and Grad-CAM analysis, for the CNN trained on the entire merged dataset to classify the task being performed. **A.** Violin plot showing the accuracy of the model. **B**. Grad-CAM results highlighting the time-channel points in the second CNN layer that contributed most to the classification. The most contributive points are in red, and the colorbar represents arbitrary units. The channels are arranged sequentially from frontal to occipital regions.

Looking at the accuracy of the model (**Figure 2.A**), we find an average classification accuracy of 75.39 ± 7.28%. Bootstrapping analysis using the binomial distribution showed that the accuracy of the model was better than the 50% chance level (95% confidence interval of the mean difference from chance: CI [19.40, 31.45], *p* < 0.0001). This result suggests that learning contextual information modified the neural correlates of scene perception allowing the network to classify the task. In order to identify which time points and EEG channels contributed most, we then looked at the Grad-CAM results (**Figure 2.B**). These results highlight that the activity of a cluster of occipitoparietal electrodes (P8/POz/O1/O2/P6/PO4/PO6/PO7/PO8/Oz) between 50 and 180ms after stimulus presentation contributed the most to task classification, a result that may reflect differences in visual stimuli processing. The same electrodes also seemed to be involved in a second section of data between 200 and 250ms. Finally, later activity appeared after 600ms, which also enabled the classification of the task, reflecting the onset of motor activity to produce the response.

In a second step, we extracted the EEG activity of participants involved in the spatial memory task (*i*.*e*., when they had to retrieve the target). We selected only trials in which participants were presented with 2 open doors and in which they successfully retrieved the position of the goal (93.47 ± 0.58% of trials per participant). This ensured that the network had sufficient trials, and used only the EEG activity associated with goal direction information as support for the classification.

The accuracy of the model (**Figure 3.A**) indicated that on average the network classified 45.41 ± 5.37 % of the tested trials correctly, performing better than the 33% expected by chance (95% CI: [3.74, 13.63], *p* = 0.0004). The Grad-CAM results (**Figure 3.B**) highlighted activity in frontal electrodes between 180 and 200ms as well as activity in occipitoparietal electrodes between 200 and 220ms as the most contributive to the goal position identification. These results suggest that early EEG activity also contains information regarding the position of the goal, irrespective of the visual features of the scene. As a control, we also performed the same analysis to detect the position of the goal but in the condition with three affordances and similarly found that the model accurately classified the position of the goal (Mean classification accuracy = 48.67 ± 6.54 %; 95% CI: [6.95, 16.89], *p* < 0.0001).

**Figure 3.**
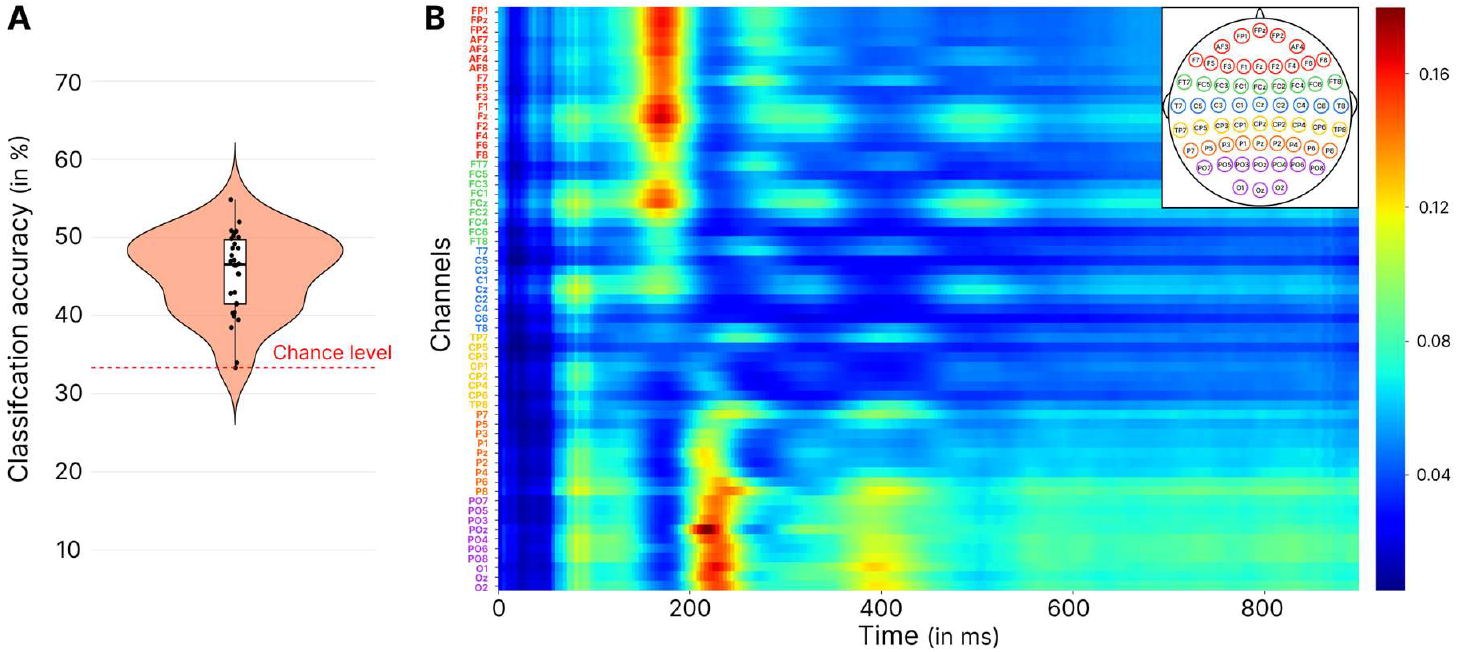
Outcomes of the EEGNet classification and Grad-CAM analysis, for the CNN trained on the dataset with the ‘two affordances’ conditions to classify the goal position. **A.** Violin plot displaying the model’s accuracy. **B**. Grad-CAM results indicating the time-channel points in the second CNN layer, with those making the highest contribution to the classification presented in red.

### Prior spatial knowledge modulates early neural activity related to affordance processing

Here, we considered the EEG activity of participants performing the scene memory task and trained the model to detect the affordances within the scene. We also tested the model on data from the spatial memory task to assess whether the information supporting decoding was modulated by learned spatial contextual information. (**Figure 4**).

**Figure 4.**
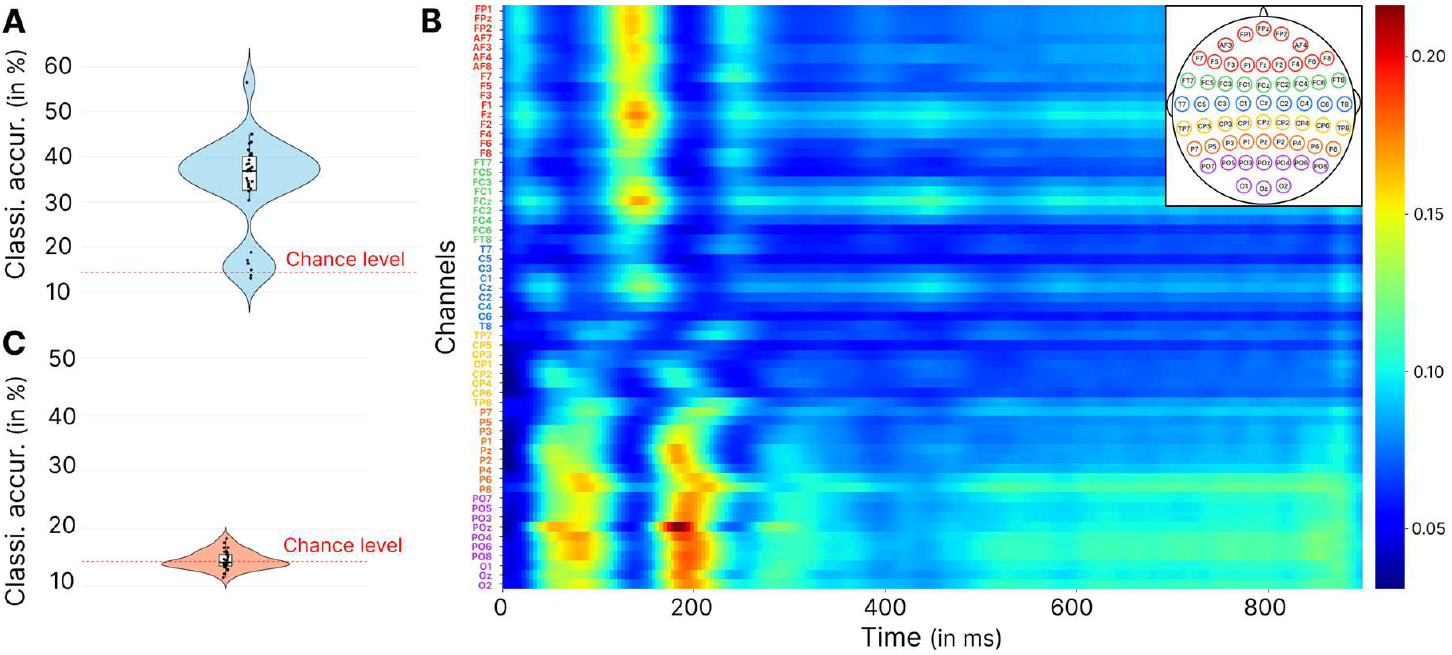
Results of the EEGNet classification and Grad-CAM analysis, for the CNN trained on the data of participants performing the scene memory task to detect the position of the affordances. **A.** Violin plot showing the accuracy of the model during the scene memory task. **B**. Grad-CAM results highlighting the time-channel points in the second CNN layer that contribute most to the classification. The most contributive points are in red, and the colorbar corresponds to arbitrary units. **C**. Violin plot of the accuracy for the model tested on the data of participants performing the spatial memory task.

The model tested on the scene memory task (**Figure 4.A**) showed an accuracy of 35.80 ± 10.45%, and bootstrap testing indicated that it performed better than the 14.29% chance level (95% CI: [10.88, 23.68], *p* < 0.0001). However, when we tested the model on EEG data from the spatial memory task (**Figure 4.C**), the classification accuracy dropped to 13.59 ± 1.78% and was no longer higher than chance level (95% CI: [-7.41, 3.11], *p* = 0.77). These results suggest that the information from neural activity enabling the classification of the position of affordances is modulated by spatial learning. The results of the Grad-CAM analysis (**Figure 4.B**) indicated that the most contributive time points for the classification were once again located in the occipitoparietal region around 200 ms, as previously suggested by modulation of the P2 component in ERP analyses (Harel et al., 2022). Interestingly, earlier occipitoparietal activity (50–100 ms) and frontal activity (150–180 ms) were also involved to a lesser extent in the classification, replicating previous findings on the neural correlates of low-visual feature processing such as the openness of visual scenes (Greene & Oliva, 2009; Hansen et al., 2018; Orima & Motoyoshi, 2023).

Finally, to test the possibility that our results could simply be explained by task differences, we conducted a supplementary analysis using a CNN trained to classify the color of the wall, another scene feature that was present in both tasks. Similar to the previous affordance processing analysis we trained a CNN on EEG data from participants performing the scene memory task and then tested it on unseen data from the same task as well as on data from the spatial memory task to assess the generalization of the information supporting classification (**Figure 5**). For the scene memory task, the CNN achieved an average accuracy of 34.34 ± 11.18%, significantly better than the 12.5% chance level (95% CI: [13.75, 25.87], *p* < 0.0001). However, when tested on data from the spatial memory task, the accuracy was on average 18.75 ± 2.09%, which was statistically higher than chance level (95% CI: [0.31, 9.65], *p* = 0.019), suggesting that the EEG information allowing the classification of wall color generalizes to some extent across tasks.

**Figure 5.**
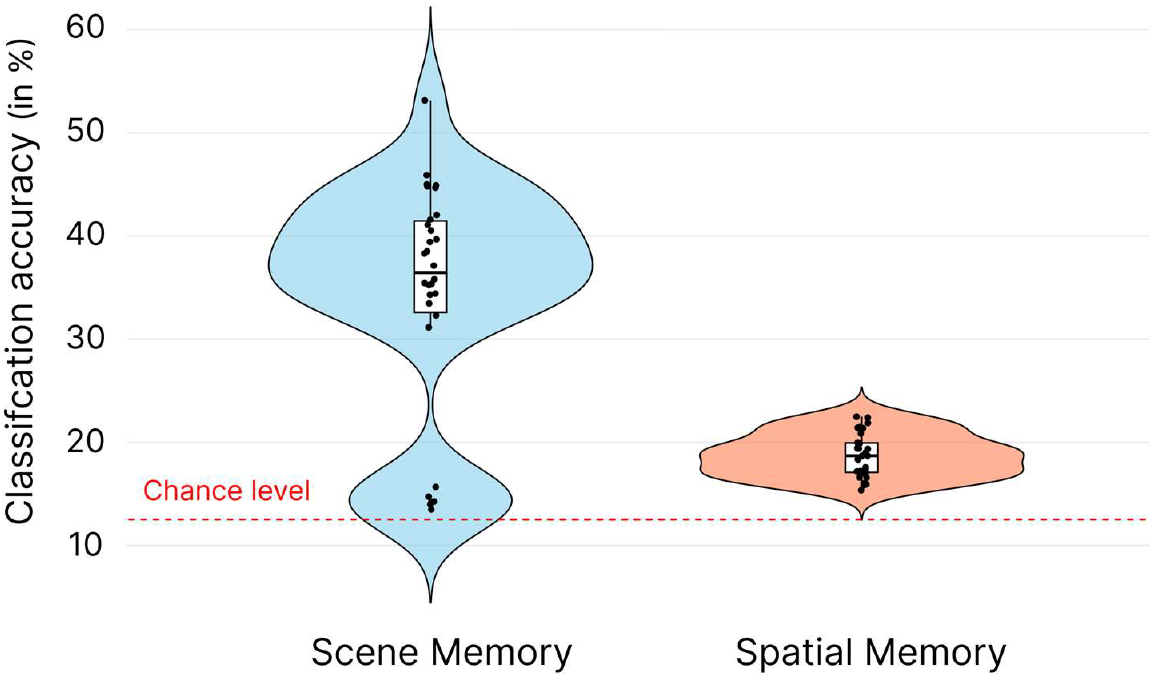
Accuracy of the CNN, trained on the data from scene memory task to detect the color of the wall (chance level at 12.5%), in the scene memory or the spatial memory task.

## Discussion

In this study, we investigated the temporal neural dynamics underlying visual perception during scene processing and its interaction with visuospatial memory information. Our results demonstrate that a CNN trained on EEG data can accurately classify several aspects of visual perception, including the color of the wall within the scene, navigational affordances, and the location of a previously learned target, primarily based on early occipitoparietal brain activity (between 100-230 ms). To further support the potential influence of top-down information suggested by this common time window of integration, we showed that a model trained on EEG data to decode affordance-related information in a scene memory task failed to generalize to the task in which participants had learned the position of a goal. In contrast, another model trained to detect wall color performed significantly above chance in the same context. Taken together, these results highlight the strong influence of top-down information related to prior spatial knowledge on early occipitoparietal neural activity during scene perception.

### Navigational affordances and visuospatial memory processing share a common early integration time window

The Grad-CAM analyses highlighted occipitoparietal activity around 200 ms as the most important contributor to the classification of navigational affordances within the scene, consistent with previous findings by Harel et al. (2022). In their study, the authors also used a multivariate approach and suggested that the location of affordances may be represented even earlier, around 130 ms. In the present study, we did not distinguish between the position and number of affordances but found that activity around both 100 ms and 200 ms contributed to the classification of affordances, with the latter time window playing a more important role. This early extraction of the position of available pathways for movement may reflect the brain’s ability to rapidly construct high-level representations of visual scenes based on categorical distinctions and global spatial properties (Orima & Motoyoshi, 2021, 2023). In support of this point, studies combining MEG measurements with deep neural network analyses have demonstrated a hierarchical progression of scene representations, integrating information from low-level features to higher-order scene features within 200 ms (Cichy et al., 2017). These findings suggest that the processing of affordances from visual scenes occurs as early as other perceptual processes, bridging the gap between scene perception (*e*.*g*., encoding the number of affordances) and action (Djebbara et al., 2019; Harel et al., 2022).

Interestingly, our results suggest that higher-level information, such as spatial contextual information about the target location, is also represented in this early occipitoparietal activity. This common integration time window is consistent with several lines of evidence for top-down influences on early visual processing during scene and object perception (Enge et al., 2023; Klink et al., 2023; Steel et al., 2021, 2023, 2024). For example, Enge et al. (2023) demonstrated that prior knowledge of an object’s function shapes early cortical markers at around 200 ms and influences higher-order visual perception (Greene & Hansen, 2020; Groen et al., 2018). Similarly, Klink et al. (2023) reported that scene familiarity modulates early EEG activity, suggesting a role for learned scene context in shaping early visual processing. The current results support this interpretation by highlighting the critical role of visuospatial memory-derived information in shaping early visual markers of scene processing. Furthermore, these results provide novel insights into a common early time window for integrating both bottom-up and top-down information, emphasizing the dynamic interplay between scene perception and prior visuospatial knowledge.

### Visuospatial memory immediately influences the early neural activity of affordance extraction

To investigate the influence of top-down modulation on early neural correlates of scene perception, we examined how prior learning of the target location affects the classification abilities of the CNN in decoding affordance information. To test this, we trained a model that accurately decoded affordances in the scene memory task, confirming that the extraction of navigational affordances is represented in early occipitoparietal activity (Dwivedi et al., 2024; Harel et al., 2022). However, this model failed to detect affordances after participants learned the path to the goal in the spatial memory task, even though the presented images were exactly the same. This suggests that the neural correlates of navigational affordances, including their number and location, are modulated by the learning of spatial contextual information about these affordances. Thus, top-down information related to prior spatial knowledge may interact with the automatic encoding of navigational affordances. This finding complements previous behavioral findings by Naveilhan et al. (2024), which showed that increasing the number of affordances decreased participants’ accuracy in the spatial memory task only. Critically, this finding does not simply reflect task differences, as highlighted by a control analysis showing significant generalization between scene memory and spatial memory tasks of a model similarly trained to decode wall colors. This lack of modulation observed for wall color features is consistent with previous findings by Hansen et al. (2018), which showed that processing of low-level visual information is rapid and minimally affected by observer-based goals. Thus, our results suggest a targeted modulation of neural activity tied to scene-specific information influenced by learning, rather than a broader, nonspecific modulation.

These findings on the interaction between scene feature integration and prior spatial information during navigational affordance processing illustrate the dynamic and complex nature of visual scene perception. From a broader perspective, this is consistent with the predictive coding framework for efficient perception (Peelen et al., 2024; Rao & Ballard, 1999) and the ecological basis of affordances (Gibson, 1979). This notion emphasizes the intrinsic connection between action and perception, highlighting the role of perception in guiding action, and is consistent with the notion of active inference, which views action and perception as integrated processes that work to minimize prediction error (Friston, 2010; Friston et al., 2017; Kaplan & Friston, 2018; Pezzulo & Cisek, 2016). Consistent with this theoretical framework, our results suggest that spatial memory content modulates the early neural correlates of navigational affordance processing to support efficient, goal-directed visual information processing. Finally, it is important to consider an alternative explanation for the modulation of affordance processing that arises from the findings of Castelhano & Witherspoon (2016). They demonstrated that knowledge of an object’s function significantly increases search efficiency by directing attention to functionally relevant areas within a scene (Bouwkamp et al., 2024). In our paradigm, participants may have explored the visual scene differently or considered other navigational options (*i*.*e*., open doors that did not lead to the goal). Future studies using eye-tracking or specific experimental manipulations, such as modifying the eccentricity of navigational affordances, could help disentangle the possibilities and shed light on this potential confounding factor.

## Conclusion

In conclusion, this work supports the notion that during scene perception neural processes associated with behaviorally relevant tasks, such as retrieving the position of a goal hidden behind a door, share a common time window of integration—approximately 100-250 ms over occipitoparietal regions—with processes representing visual information, such as navigational affordances. The present results also demonstrate an immediate influence of the newly learned spatial knowledge on the early neural activity associated with scene processing, modulating even the first stages of visual scene perception. Thus, knowing where to go may shape what you see.

## Data availability

The complete set of data, stimuli generated and code used for preprocessing and CNN is available on the OSF repository : https://osf.io/5cqxy/?view_only=2957dac1a1304c2db6e1fb3b056c5008.

## Author Contributions

**Clément Naveilhan**: Conceptualization; Formal analysis; Investigation; Methodology; Writing—Original draft; Writing—Review & editing. **Raphaël Zory:** Funding acquisition; Writing—Review & editing. **Stephen Ramanoël**: Conceptualization; Funding Acquisition; Methodology; Project administration; Supervision, Writing—Review & editing.

## Acknowledgements

This research was made possible by the generous participation of volunteer participants, to whom the authors are sincerely grateful. We also thank Catherine Buchanan for her careful reading of the manuscript and her feedback.

This work was supported by the French government through the France 2030 investment plan managed by the National Research Agency (ANR), as part of the Initiative of Excellence Université Côte d’Azur under reference number ANR-15-IDEX-01 and, in particular, by the interdisciplinary Institute for Modeling in Neuroscience and Cognition (NeuroMod) of Université Côte d’Azur.

